# Integrated modeling of peptide digestion and detection for the prediction of proteotypic peptides in targeted proteomics

**DOI:** 10.1101/399436

**Authors:** Zhiqiang Gao, Cheng Chang, Yunping Zhu, Yan Fu

**Affiliations:** NCMIS, RCSDS, Academy of Mathematics and Systems Science, Chinese Academy of Sciences, Beijing 100190, China; State Key Laboratory of Proteomics, Beijing Proteome Research Center, National Center for Protein Sciences (Beijing), Beijing Institute of Lifeomics, Beijing 102206, China; School of Mathematical Sciences, University of Chinese Academy of Sciences, Beijing 100049, China

## Abstract

**Motivation:** The selection of proteotypic peptides, i.e., detectable unique representatives of proteins of interest, is a key step in targeted shotgun proteomics. To date, much effort has been made to predict proteotypic peptides in the absence of mass spectrometry data. However, the performance of existing tools is still unsatisfactory. One crucial reason is their neglect of the close relationship between protein proteolytic digestion and peptide detection.

**Results:** We present an algorithm (named AP3) that firstly considers peptide digestion probability as a feature for proteotypic peptide prediction and demonstrated peptide digestion probability is the most important feature for accurate prediction of proteotypic peptides. AP3 showed higher accuracy than existing tools and accurately predicted the proteotypic peptides for a targeted proteomics assay, showing its great potential for assisting the design of targeted proteomics experiments.

**Availability and Implementation:** Freely available at http://fugroup.amss.ac.cn/software/AP3/AP3.html.

**Supplementary Information:** Supplementary data are available at *Bioinformatics* online.

## Introduction

Targeted proteomic assays, such as multiple reaction monitoring (MRM) experiments, are capable of the sensitive identification and quantification of proteins of interest and have become a promising powerful tool for biological or clinical studies, such as the verification of candidate biomarkers (Parker and Borchers, 2014; Rifai *et al.*, 2006). The first key step in developing an MRM assay is the selection of proteotypic peptides, i.e., peptides that are unique representatives of their corresponding proteins and possess good mass spectrometry (MS) detectability (Demeure *et al.*, 2014). There are two major approaches for proteotypic peptide selection, i.e., the empirical approach and the computational approach. The former selects previously identified peptides as proteotypic peptides, while the latter predicts proteotypic peptides from the physiochemical properties of peptides. Although the empirical approach has been successfully applied in MRM-MS assays, it has some limitations. For example, not all target proteins have experimental evidence, especially proteins identified by literature mining. Moreover, there is randomness in peptide detection, i.e., some peptides identified in one experiment may not be identified in the next experiment. Thus, researchers are paying increased attention to the computational approach. However, the mechanisms underlying peptide detection are still unclear, which hinders the development of accurate algorithms to predict proteotypic peptides.

To date, much effort has been devoted to understanding the mechanisms underlying peptide detection. In early research, Le et al. (Le Bihan *et al.*, 2004) and Ethier et al. (Ethier and Figeys, 2005) proposed empirical score functions based on hydrophobicity, peptide length and isoelectric point. Recently, several algorithms (Fusaro *et al.*, 2009; Webb-Robertson *et al.*, 2008; Mallick *et al.*, 2007; Sanders *et al.*, 2007; Wedge *et al.*, 2007; Eyers *et al.*, 2011; Qeli *et al.*, 2014) have been developed to predict proteotypic peptides or high-responding peptides based on supervised machine learning. In such algorithms, designing the features that describe peptides is a key problem. Many factors govern the likelihood of observing a peptide in a proteomics experiment, such as the physicochemical properties of the peptide, the abundance of the associated protein, and the identification procedure (Tang *et al.*, 2006; Jarnuczak *et al.*, 2016). Tang et al. (Tang *et al.*, 2006) proposed the concept of peptide detectability, which was defined as the probability that a peptide would be observed in a standard sample analyzed by a standard proteomics routine. They also invented a machine learning algorithm based on 175 features derived from the peptide sequence to predict peptide detectability. Later, Sander et al. (Sanders *et al.*, 2007), Mallick et al. (Mallick *et al.*, 2007) and Eyers et al. (Eyers *et al.*, 2011) developed algorithms using 596, 1010, and 1186 features, respectively. These features included mainly AAindex (Kawashima *et al.*, 1999)-derived features and other sequence-derived features. Recently, Muntel et al. (Muntel *et al.*, 2015) considered protein abundance as an additional feature and obtained improved performance. However, information on protein abundance is generally unavailable in the absence of experimental MS data.

A key limitation of the above algorithms is that they do not make (full) use of the protein proteotypic digestion information. As we know, a typical bottom-up proteomics experiment can be divided into two continuous processes: protein proteolytic digestion and peptide detection. An easily neglected fact is that the products of protein proteolytic digestion are uncertain. That is, we do not know exactly which peptides and what proportions of them will be produced by digestion and undergo subsequent MS detection. Therefore, the accurate prediction of peptide detectability requires considering the process of protein proteolytic digestion. Previous studies have demonstrated that the commonly used enzyme trypsin exclusively cleaves the C-terminal to arginine or lysine (Olsen *et al.*, 2004), and this process is always incomplete (Siepen *et al.*, 2007). In addition to the local conformation, tertiary structure and experimental condition, the cleavage probability of a tryptic site is mainly influenced by the amino acids surrounding the site (Siepen *et al.*, 2007). Several algorithms have been proposed to predict the cleavage probabilities of tryptic sites from the adjacent amino acids (Lawless and Hubbard, 2012; Fannes *et al.*, 2013).

Here, we propose a new machine learning algorithm named AP3 (short for Advanced Proteotypic Peptide Predictor) that, for the first time, integrates the peptide digestion and detection processes together to predict proteotypic peptides. Specifically, it incorporates the peptide digestion probability predicted in the first stage into the peptide detectability prediction model in the second stage. Training on a yeast dataset showed that peptide digestion probability was the most important feature for predicting proteotypic peptides, increasing the 10-fold cross-validation accuracy (area under the ROC curve, AUC) from 0.8891 to 0.9428. Furthermore, we demonstrated that the AP3 model trained on the yeast dataset retained good predictive power over three independent comprehensive datasets, including *E. coli* (AUC 0.9383), mouse (AUC 0.9311) and human (AUC 0.9252), and exhibited superior performance (15.3%-17.5% higher in AUCs) to that of existing tools, i.e., PeptideSieve (Mallick *et al.*, 2007), CONSeQuence (Eyers *et al.*, 2011), ESP Predictor (Fusaro *et al.*, 2009), and PPA (Muntel *et al.*, 2015). Lastly, we showed that AP3 can be used to effectively select proteotypic peptides for MRM-MS assay development in the absence of experimental MS data.

## Algorithm

As **Fig. 1** illustrates, AP3 has two major components: a digestion probability predictor and a peptide detectability predictor. A cleavage model is trained first to predict the cleavage probabilities of tryptic sites. Then, the peptide digestion probability is calculated and integrated into the peptide detectability predictor as a feature that characterizes peptides. Finally, feature selection is performed, and the peptide detectability prediction model is trained using the selected features.

**Figure 1.**
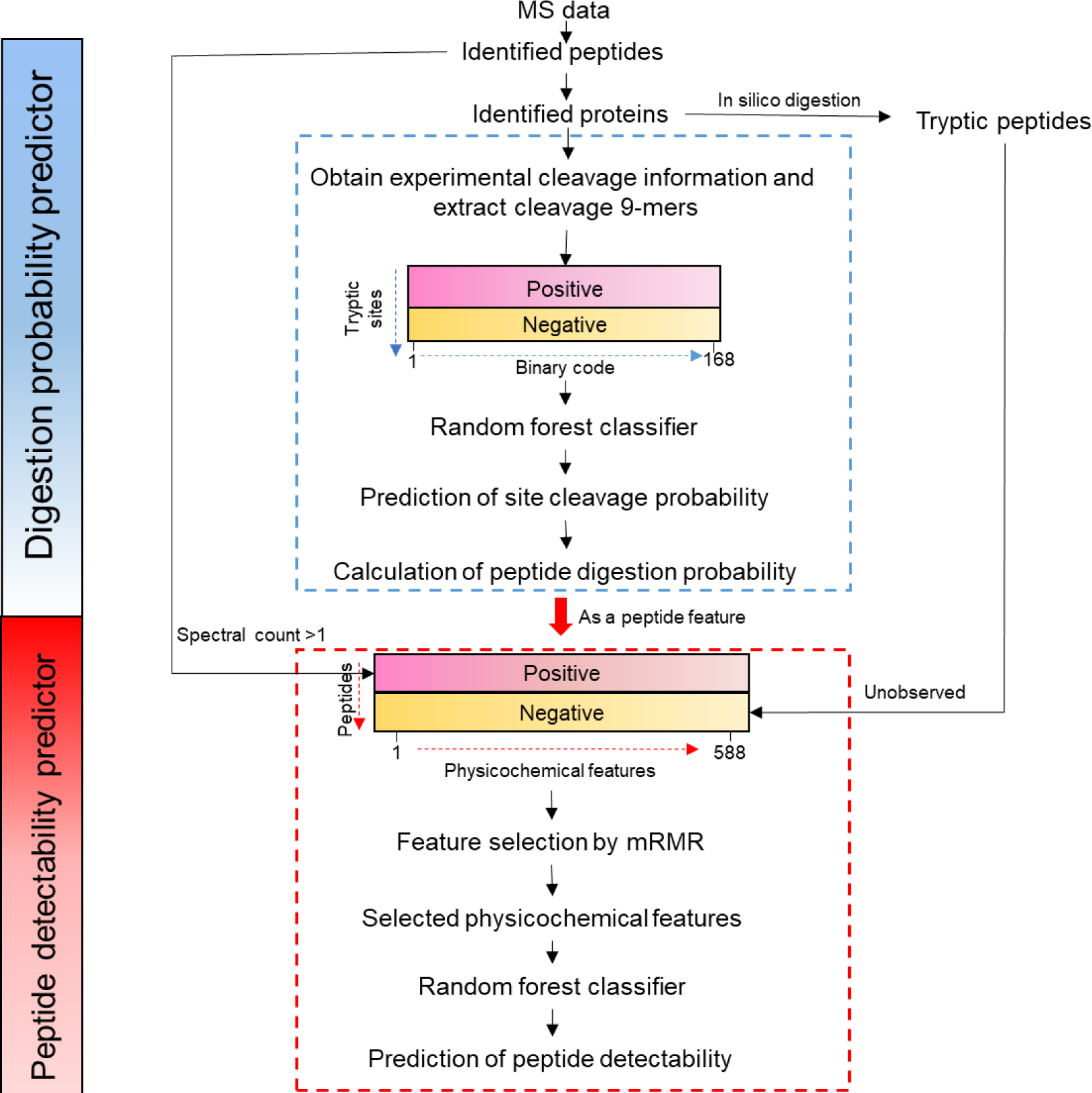
Overview of the development of the AP3 algorithm.

### Peptide digestion probability predictor

#### Constructing the training set

Identified peptides are mapped to their corresponding protein sequences. The cleavage information of tryptic sites in the identified protein sequences is collected, including the spectral count (SC) of the peptides observed on the left of the tryptic site (L), the SC of the peptides observed on the right of the tryptic site (R) and the SC of the peptides in which this tryptic site is a missed cleavage (O). Then, tryptic sites are classified as positive sites if (1) L is at least 1, (2) R is at least 1 and (3) O is zero and as negative sites if both L and R are zero and O is at least 2. Previous studies have demonstrated that the cleavage probability of a tryptic site is influenced mainly by the amino acids adjacent to the tryptic site (Hubbard *et al*., 1998; Fannes *et al.*, 2013). Therefore, for each positive or negative site, a 9-mer is taken consisting of the tryptic site and four residues on both sides. If the tryptic site is located on the N or C terminus of a protein, resulting in insufficient amino acids to form a 9-mer, the character Z is added to make up the 9-mer. Each character in the 9-mer, except for the tryptic site (arginine or lysine), is converted into a 21-dimensional binary vector that indicates whether one of the 20 amino acids or the character Z appears. For one binary vector, if one amino acid appears, the corresponding position is 1, and other positions are set to 0. Thus, each 9-mer is converted into one 168-dimensional binary vector that retains both the position and residue-specific information. The positive sites are labeled with 1, and the negative sites are labeled with 0.

#### Random forest classifier

A random forest is a nonlinear ensemble classifier consisting of a collection of independent unpruned trees (Breiman, 2001). The randomness of this algorithm is reflected in two aspects: randomly selecting a training subset for each tree by bootstrap and randomly selecting features for the best split at each node. The parameters setting of random forest were described in the **Supplementary Note, Section 1**.

#### Calculating peptide digestion probability

For the remaining proteins, we perform an in silico digestion with a dataset-dependent peptide length range and up to 2 missed cleavage sites and then predict the cleavage probabilities of all tryptic sites associated with the digested peptides using the trained cleavage probability model. The dataset-dependent peptide length range means that the longest/shortest digested peptide length allowed is set to the longest/shortest length of identified peptides in the dataset, respectively. The digestion probability of a peptide, which is defined as the probability of the peptide being produced by the protein digestion process, is calculated according to the following formula:

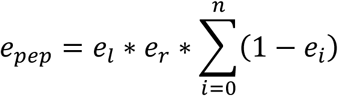

where *e*_*pep*_ is the digestion probability of this peptide, *e*_*l*_ and *e*_*r*_ are the cleavage probabilities of the left and right tryptic sites of the peptide sequence, respectively, e_*i*_ is the cleavage probability of the missed cleavage site *i* in the peptide sequence, and *n* is the number of missed cleavage sites in the peptide sequence.

### Peptide detectability predictor

#### Constructing the training set

In the data preprocessing phase, we filtered proteins according to SC and sequence coverage to ensure high confidence in the remaining proteins. All the digested peptides of the remaining proteins theoretically have the chance of being observed. Identified peptides with SC larger than 1 are taken as positive peptides, and unobserved digested peptides are taken as negative ones.

To date, the mechanism underlying peptide detection is still not clear, so we collect as many related computer features as possible. Finally, a diverse set of 588 features are used to characterize each peptide **(Supplementary Note, Section 2; Supplementary Table 1; Supplementary Fig. 1)**. It is notable that we add the peptide digestion probability to the feature set of the peptide detectability model for the first time.

#### Feature selection and random forest classifier

The minimum redundancy maximum relevance (mRMR) approach (Ding and Peng, 2003) is used for feature selection **(Supplementary Note, Section 3; Supplementary Fig. 2)**, then random forest is used to model the peptide detectability with selected features. During the construction of the training set of peptide detectability model, the number of identified peptides is far less than the number of unobserved digested peptides in most cases. Thus, to avoid this imbalance problem of the training set, we employed the down-sampling technique (Chen *et al.*, 2004) for the negative peptides (i.e., the majority class). In brief, the number of training samples for each class is set to the size of the minority class, and samples within the majority class are selected randomly together with the minority class to form a balanced training set. Then, since the scales of the selected features are different, z-score normalization (Jain *et al.*, 2005) is employed for each feature to obtain a zero mean and unity variance. A 10-fold cross-validation AUC is calculated to evaluate the performance of our model. The trained random forest model can be used to predict the detectability of each peptide. Finally, we sort the peptides for each protein of interest in descending order by the peptide detectability and select the top peptide as the proteotypic peptide of this protein.

## Results

### Datasets

Here, a public large-scale yeast dataset (Hebert *et al.*, 2014) was used as training data for our algorithm AP3. To validate the generalization performance of our algorithm, we also used three publicly available datasets from other organisms: *E. coli* (Schmidt *et al.*, 2016), mouse (Malmström *et al.*, 2016) and human (Wilhelm *et al.*, 2014). For the human dataset, the data of the lymph node and salivary gland were used. The raw files from the four public datasets were downloaded and reanalyzed. Table 1 provides a summary of the four public datasets used in our study. The details of MS data preprocessing were described in **Supplementary Note, Section 4.**

**Table 1.** Description of the four large-scale public datasets used for the development and evaluation of AP3

| Dataset | yeast | human | *E. coli* | mouse |
| --- | --- | --- | --- | --- |
| Reference | ref 1 | ref 2 | ref 3 | ref 4 |
| Species | *Saccharomyces cerevisiae* | *Homo sapiens* | *Escherichia coli* | *Mus musculus* |
| Instrument | Orbitrap Fusion | LTQ Orbitrap | LTQ Orbitrap Elite | Q Exactive Plus |
| Protein groups observed | 3959 | 6422 | 2898 | 5304 |
| Peptides observed | 43088 | 80490 | 38035 | 58184 |

References 1-4:
1. Hebert, A.S. et al. (2014) The One Hour Yeast Proteome. *Mol. Cell. Proteomics*, 13, 339–347. 2. Wilhelm, M. et al. (2014) Mass-spectrometry-based draft of the human proteome. *Nature*, 509, 582–587. 3. Schmidt, A. et al. (2016) The quantitative and condition-dependent *Escherichia coli* proteome. *Nat. Biotechnol.*, 34, 104–110. 4. Malmström, E. et al. (2016) Large-scale inference of protein tissue origin in gram-positive sepsis plasma using quantitative targeted proteomics. *Nat. Commun.*, 7, 10261.

### Performance of peptide digestion probability predictor

There were 3959 proteins with 43088 peptides identified in the yeast dataset. After filtering by the sequence coverages and SCs of proteins, 1556 proteins with 29171 peptides remained. Following the construction rules of the cleavage probability training set described in the Methods section, 7778 positive tryptic sites and 4854 negative tryptic sites were obtained. As shown in **Supplementary Fig. 3**, the trained cleavage probability model had a 10-fold cross-validation AUC of 0.9754, and the average AUC for the three test datasets was 0.9747. These results indicated that the cleavage probability predictor could accurately predict the cleavage probabilities of tryptic sites.

### Feature selection for peptide detectability prediction

According to the down sampling technique, 25363 identified peptides with at least 2 SCs were taken as positive peptides, and 25363 peptides that were randomly selected from the unobserved digested peptides were taken as negative peptides. For each peptide, 588 features were calculated, including the peptide digestion probability. Using the feature selection algorithm mRMR, 31 features were ultimately selected with the maximum AUC (**Fig. 2 and Table 2**). We grouped the 31 selected features into six categories: digestion, hydrophobicity, structural, charge, energy and other. There was broad agreement that the hydrophobicity, structure, charge and energy terms had a large impact on the peptide detection (Fusaro *et al.*, 2009; Webb-Robertson *et al.*, 2008; Mallick *et al.*, 2007; Sanders *et al.*, 2007; Eyers *et al.*, 2011). Hydrophobicity was represented by the hydrophobic residues and hydrophobicity coefficient in reversed-phase high-performance liquid chromatography (RP-HPLC). Nine of the 31 selected features were related to secondary or tertiary structures, suggesting that peptide structure also influenced peptide detection. Charge played an important role in peptide detection, which is consistent with (Mallick *et al.*, 2007). The selected feature “Activation Gibbs energy of unfolding” was consistent with a previous study, which claimed that Gibbs free-energy transfer between amino acids led to an increased response in peptides with nonpolar regions (Cech and Enke, 2000).

**Figure 2.**
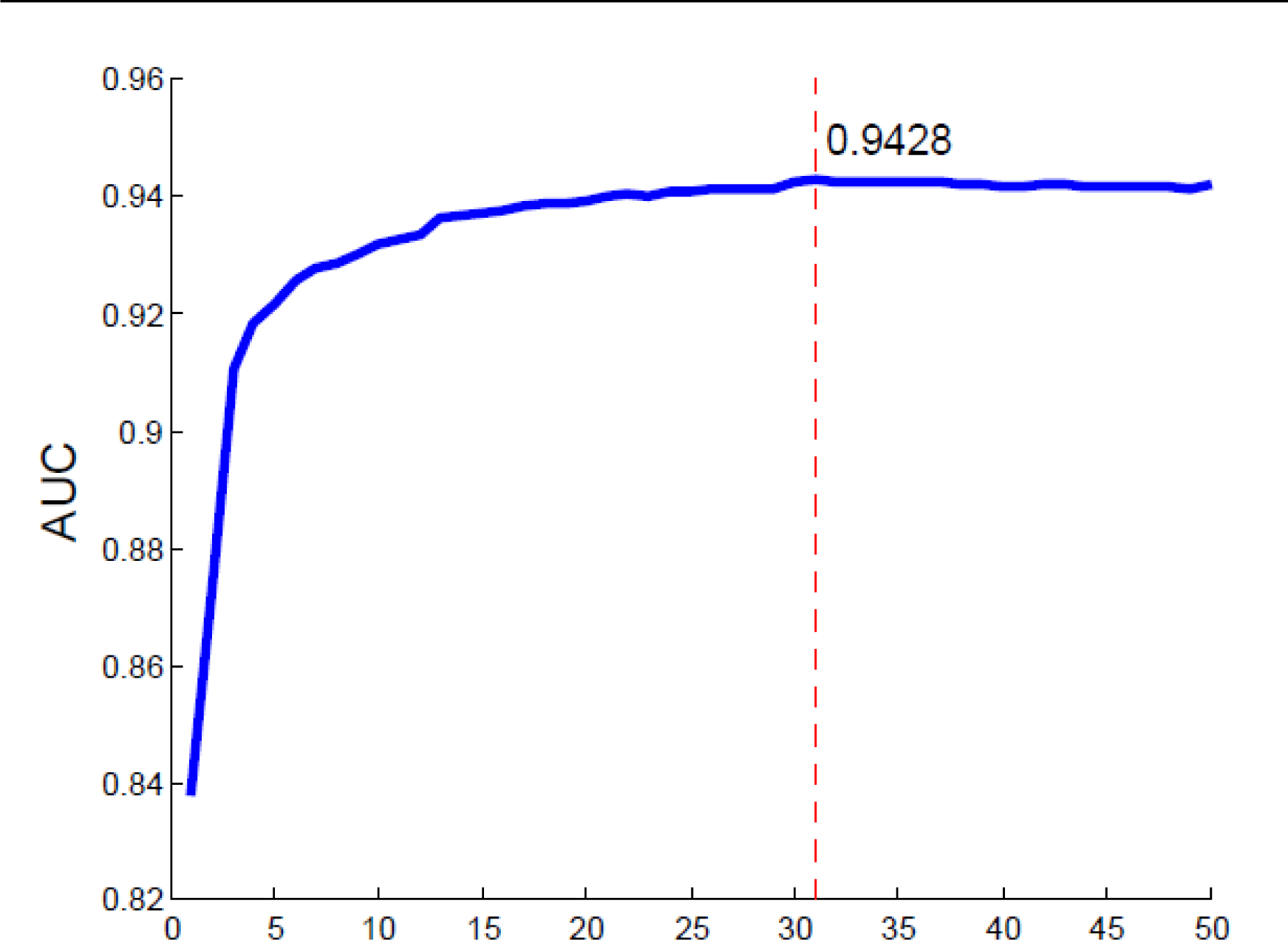
The incremental feature selection (IFS) curve was plotted by 10-fold cross-validation, as the top 50 features selected by mRMR were added successively to the random forest model. Ultimately, we selected 31 features with a maximum AUC of 0.9428.

**Table 2.**
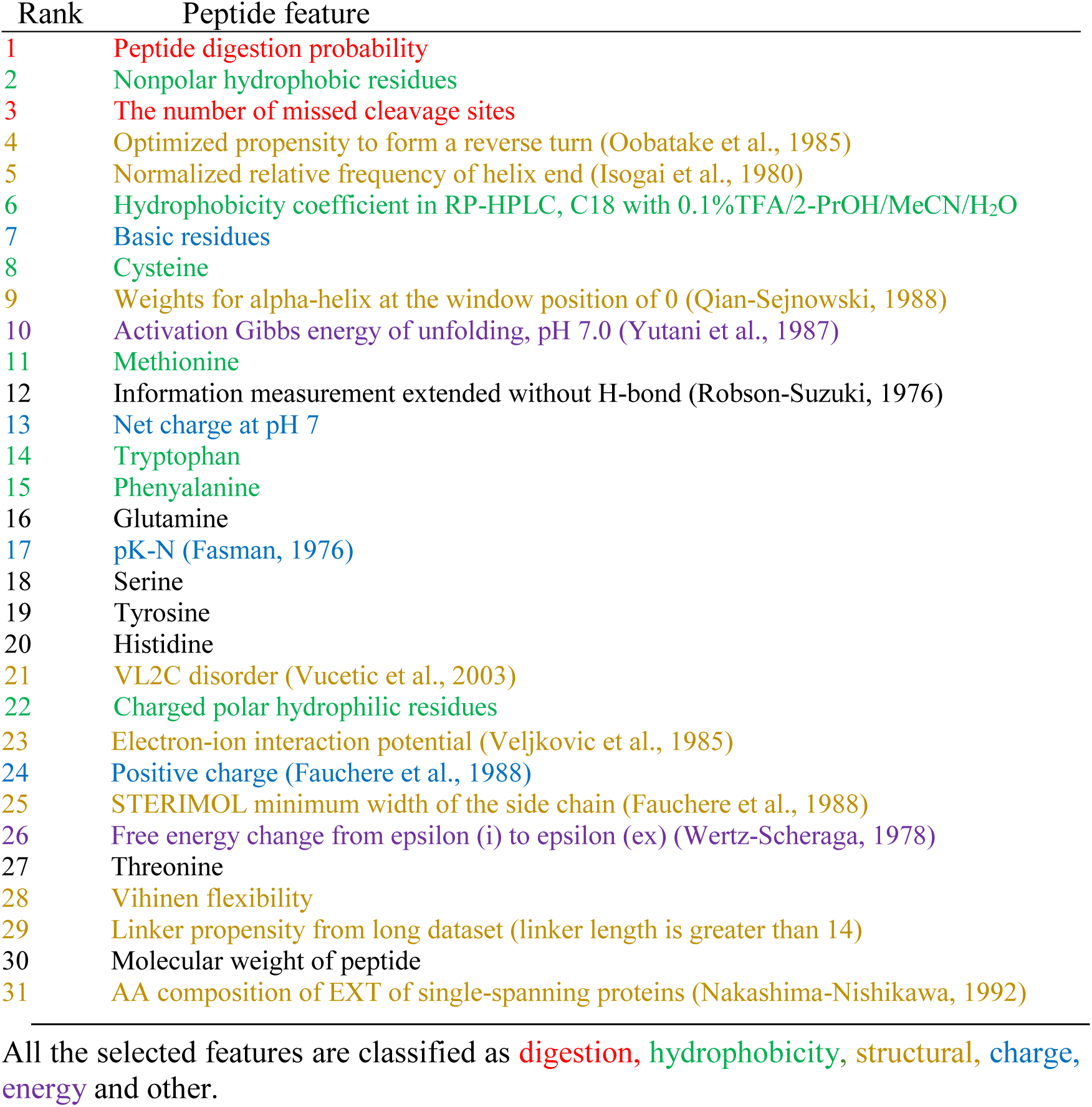
Selected features of the yeast dataset obtained using the mRMR method

To validate the feature selection performance, we compared the prediction performances using all 588 features and the 31 selected features, respectively. The results indicated that the simplified model using the 31 selected features had similar accuracy to that of the full model using all 588 features and showed better generalization ability than the full model (**Fig. 3**).

**Figure 3.**
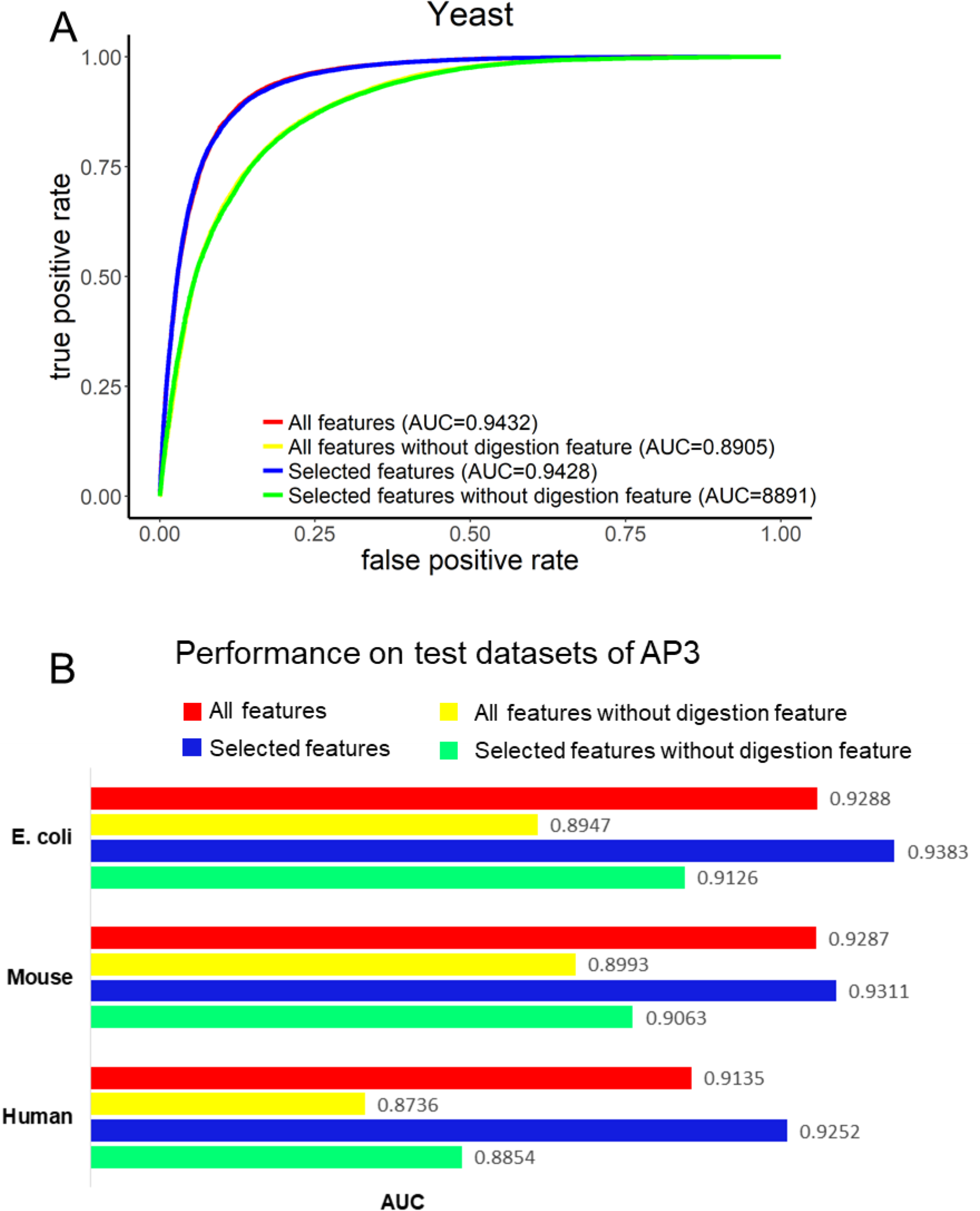
The performance of AP3. (A) The 10-fold cross-validation ROC curves were plotted for the peptide detectability model on the training set for four cases: using the digestion feature (peptide digestion probability) and other 587 features (red), using other 587 features without the digestion feature (yellow), using the digestion feature together and other 30 selected features (blue), using other 30 selected features without the digestion feature (green). (B) Comparison of the generalization abilities of all features-based model (red), all features without digestion feature-based model (yellow), selected features-based model (blue) and selected features without digestion feature-based model (green) on three independent test datasets.

Notably, our proposed feature peptide digestion probability had good performance in the feature selection process. First, we found the peptide digestion probability was the top feature among the selected features (**Table 2**), and this conclusion had also been validated on the three test datasets (**Supplementary Table 2**). Second, we compared the generalization ability of peptide detection algorithms before and after the feature was included in the selected features on the three test datasets. By incorporating peptide digestion probability, the 10-fold cross-validation AUC increased 6.0% on the training dataset (**Fig. 3A**), and the AUCs increased 2.8%, 2.7% and 4.5% on the three test datasets (*E. coli*, mouse, human), respectively (**Fig. 3B**). Second, we calculated the detectability score for each proteotypic and undetected digested peptide for the three test datasets with/without the feature peptide digestion probability (**Supplementary Fig. 4**). The different distributions between the peptide digestion probabilities of the proteotypic and undetected peptides and the larger scores for the proteotypic peptides indicated that our model was able to classify the proteotypic and undetected peptides. Importantly, 91.3% of identified peptides received scores above 0.5 averagely on three test datasets, which indicated that the majority of identified peptides could be predicted correctly. The above conclusions based on the selected features were also confirmed by comparing the generalization ability of peptide detection algorithms before and after the peptide digestion probability was included in all features on three test datasets (**Fig. 3B, Supplementary Fig. 4**). Third, using only one feature peptide digestion probability, the random forest classifier exhibited a 10-fold cross-validation AUC of 0.8552 **(Supplementary Fig. 5A)**. The above analyses illustrated that peptide digestion probability was the most important feature for predicting proteotypic peptides.

### Relationship between peptide digestion probability and the number of missed cleavage sites

Integrating peptide digestion probability into the feature set of the peptide detectability model was inspired by the close relationship between protein proteolytic digestion and peptide detection and the fact that peptide digestion probability directly reflects the degree of protein proteolytic digestion. However, other features may also have a relationship with peptide digestion, such as the number of missed cleavage sites in the peptide sequence. To demonstrate that peptide digestion probability is a better representative of peptide digestion than the number of missed cleavage sites for peptide detection, we performed a comparative analysis of the two features. First, each of the two features was combined with other 586 features to form two sets of 587 features. The two feature sets resulted in 10-fold cross-validation AUCs of 0.9421 (the feature set including peptide digestion probability) and 0.8910 (the feature set including missed cleavage site number), respectively. We then generated three models using the two features separately and together. As shown in **Supplementary Fig. 5A**, the 10-fold cross-validation AUC of the model using both features was 0.8665, the 10-fold cross-validation AUC of the model using only peptide digestion probability was 0.8552, and the 10-fold cross-validation AUC of the model using only the number of missed cleavage sites was 0.8076. Second, the KL distance was calculated to measure the distinguishing ability of peptide digestion probability and missed cleavage number for proteotypic peptides and unobserved digested peptides. The results showed that the KL distance of peptide digestion probability (3.09) was larger than that of the number of missed cleavage sites (1.66). Third, the peptide digestion probability distributions of peptides with different numbers of missed cleavage sites showed that the more missed cleavage sites the peptides had, the smaller the peptide digestion probability was (**Supplementary Fig. 5B**). All the analysis results demonstrated that peptide digestion probability was a more powerful feature for peptide detectability prediction than the number of missed cleavage sites although the two features have negative correlations.

### Performance of peptide detectability predictor

The peptide detectability prediction model with selected features obtained a 10-fold cross-validation AUC of 0.9428 on the yeast training dataset (**Fig. 3A**). Three independent public datasets (*E. coli,* mouse and human) were used to measure the generalization performance. The AUCs for the three independent test datasets (*E. coli,* mouse and human) were 0.9383, 0.9311, and 0.9252, respectively (**Fig. 3B**). These results illustrated that our model retained good predictive power on datasets in other organisms.

To evaluate the performance of our algorithm, we compared AP3 with existing available tools, including PeptideSieve (Mallick *et al.*, 2007), CONSeQuence (Eyers *et al.*, 2011), ESP Predictor (Fusaro *et al.*, 2009), and PPA (Muntel *et al.*, 2015) **(Supplementary Note, Section 5)**. The ROCs of different peptide detectability tools on the three test datasets are shown in **Fig. 4**. The results showed that AP3 outperformed other tools with respect to the true positive rate at any given false positive rate. On three test datasets, AP3 exhibited superior performance: average increases of 9.0%, 14.5%, 25.2% and 35.6% in AUC compared with the four available tools, PeptideSieve, ConSequence, PPA and ESP Predictor, respectively.

**Figure 4.**
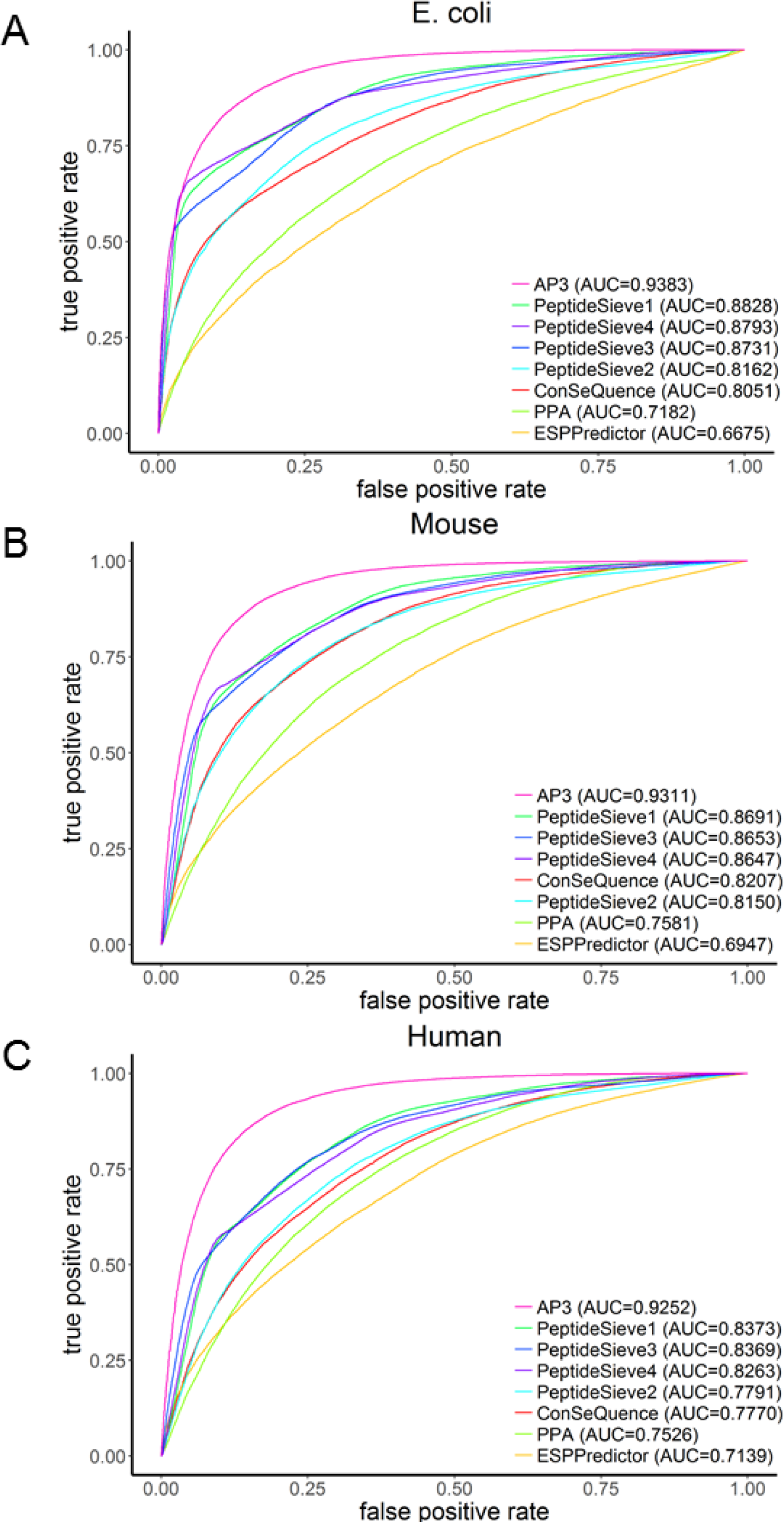
Performance comparison between AP3 and other tools on three independent test datasets. Abbreviations: PeptideSieve1: PeptideSieve_ICAT_ESI; PeptideSieve2: PeptideSieve_MUDPIT_ESI; PeptideSieve3: PeptideSieve_PAGE_ESI; and PeptideSieve4: PeptideSieve_PAGE_MALDI.

### MRM assay validation

One direct application of a peptide detectability algorithm is selecting proteotypic peptides for targeted proteomics assays. We applied AP3 and other peptide detectability tools to an MRM dataset published by Fusaro et al. (Fusaro *et al.*, 2009). Briefly, this dataset consists of 14 proteins, each of which has several experimentally validated MRM peptides. We first predicted the peptide detectability of all digested peptides of these proteins, and then the protein sensitivity, which was the percentage of proteins with one or more peptides predicted by the predictor to be among the five highest responding peptides, was used to measure the performance (Fusaro *et al.*, 2009). The protein sensitivity of AP3 was 100% (14/14), while the protein sensitivities of ESP Predictor, PPA, and PeptideSieve were all 93% (13/14) (**Supplementary Table 3**). The results indicated that AP3 was capable of accurately selecting proteotypic peptides for MRM-MS assays.

## Conclusions and discussions

In this study, we proposed an algorithm, named AP3, for predicting peptide detectability. For the first time, we integrate the peptide digestion probability into the feature set of the peptide detectability model as a novel and significant feature. We demonstrate that incorporating peptide digestion probability can significantly increase the performance of the peptide detectability model. Another advantage of AP3 is its ease of use. AP3 only needs the sequences of proteins of interest as input and provides a user-friendly graphic-user interface, which facilitates its usage. It enables the selection of candidate proteotypic peptides for proteins of interest in the absence of high-quality MS-based experimental evidence, especially for proteins identified by methods other than proteomics, such as genomic experiments or literature mining.

In summary, AP3 is a robust algorithm for the prediction of proteotypic peptides for a given protein based entirely on the peptide sequence and its neighboring regions in its parent protein. At the same time, we propose and demonstrate that peptide digestion probability is the most important feature for peptide detectability prediction. This study may have a significant effect on improving protein quantification, designing targeted proteomics assays, and developing biological biomarkers for early diagnosis and therapy.

## Supporting information

## Acknowledgments

This work was supported by the Strategic Priority Research Program of CAS (XDB13040600), the National Basic Research Program of China (2017YFA0505002 and 2015CB554406), the National Natural Science Foundation of China (21605159 and 21475150), the Innovation Program (16CXZ027), the International S&T Cooperation Program of China (2014DFB30010), and the NCMIS CAS.

## Author contributions

Y.F, C.C. and Y.Z. designed the algorithms and experiments. Z.G. and C.C. implemented the algorithms and performed the data analysis. Z.G. and C.C. wrote the initial manuscript. All authors edited and approved the final manuscript.

