## Supplementary material for "Integrated modeling of peptide digestion and detection for the prediction of proteotypic peptides in targeted proteomics"

to

|| Contributed equally to this work

* To whom correspondence should be addressed

### Supplementary Notes

#### Section 1: The parameters setting of Random forest

Generally, as the number of trees increases, the generalization error of the forest would decrease and converge gradually to a limit, but the running time would also increase rapidly (Breiman, 2001). Based on our test, using 200 trees achieves a good trade-off between accuracy and efficiency for our problem (data not shown). The number of randomly selected features for each node is set as its default value, that is, the square root of the number of all features.

#### Section 2: Characterizing each peptide using 588 features

The peptide length, number of missed cleavage sites, peptide molecular weight and frequencies of 20 amino acids are first calculated according to the peptide sequence. Then, 544 amino acid-related physicochemical features from AAindex (Kawashima *et al.*, 1999) are considered. For each of the 544 AAindex features, the numerical values of constituent amino acids in a peptide are averaged to produce a single value. Then, 20 additional features collected from previous studies (Webb-Robertson *et al.*, 2008; Eyers *et al.*, 2011; Tang *et al.*, 2006; Braisted *et al.*, 2008; Vucetic *et al.*, 2003) about peptide detection are added. Finally, we find that the peptide digestion probability can distinguish proteotypic peptides from unobserved digested peptides (KL distance = 3.09, **Supplementary Fig. 1**). The KL distance is a measure of the "distance" between two distributions. A larger KL distance indicates that the feature distribution of positive peptides is strongly different from the feature distribution of negative peptides. Thus, we add the peptide digestion probability to the feature set of the peptide detectability model for the first time.

#### Section 3: Feature selection on peptide detectability training set

To characterize the behavior of peptides in MS, 588 physicochemical features for each peptide are calculated. However, some features may be highly correlated with others. For example, peptide length and molecular weight have similar distributions in the positive set and the negative set (**Supplementary Fig. 2**). Therefore, feature selection is performed by using the minimum redundancy maximum relevance (mRMR) approach (Ding and Peng, 2003). mRMR can sort features by considering the relevance to the dependent variable and the redundancy with higher-ranked features. To select the minimum set of features with sufficient predictive performance, the top 50 features are added into the peptide detectability model one by one in order. A 10-fold cross-validation strategy is adopted to evaluate the performance of the trained model as the number of added features increases.

#### Section 4: MS data preprocessing

For the raw files produced by liquid chromatography coupled with tandem MS, peaks are searched against the corresponding organism sequences in the UniProt database using the Andromeda search engine included in the software MaxQuant (Cox and Mann, 2008) (version 1.6.0.1). Carbamidomethylation on cysteine is set as a fixed modification. Oxidation on methionine and protein N-terminal acetylation are set as variable modifications. Peptides are searched using fully tryptic cleavage constraints, and up to two missed cleavage sites are allowed. The precursor mass tolerance is set to 20 ppm for the first search (for the identification of the maximum number of peptides for mass and retention time calibration) and 4.5 ppm for the main search (for the refinement of the identifications). The mass tolerance for fragment ions is set to 0.05

Da for Q Exactive Plus and 0.5 Da for other instruments. False discovery rates at the protein and peptide levels are both set to 1%.

To construct a more precise training set, identified proteins are filtered to ensure that the remaining proteins are highly confident. We define the SC of a protein as the sum of its related peptide SCs. It is reasonable to assume that the larger the SC or sequence coverage of a protein, the higher is its confident. Therefore, proteins are sorted twice according to their SCs and sequence coverages respectively. Proteins that appear in the top 50% of both ranks remain for further analysis.

#### Section 5: Evaluating the performance of other peptide detectability tools

PeptideSieve, CONSeQuence and ESP Predictor were all trained on a yeast dataset, while PPA was trained on a human dataset. PeptideSieve took protein sequences as the input in the FASTA format. The maximum number of missed cleavage sites was set to 2, and the maximum peptide mass was set to 6000 Da. We tested all types of PeptideSieve (PAGE_ESI, PAGE_MALDI, MUDPIT_ESI, MUDPIT_ICAT). CONSeQuence was run online (http://king.smith.man.ac.uk/CONSeQuence/) with the number of internal missed cleavage sites set to 2 and the prediction type set to ANN only. The ESP Predictor was also run online (https://genepattern.broadinstitute.org/) using the default parameters. The Perl script PPA.pl was downloaded from the website http://software.steenlab.org/rc4/PPA.php. We provided a file containing peptide sequences as PPA.pl and obtained an output file that contained peptide detectability for every peptide.

#### Supplementary Figures

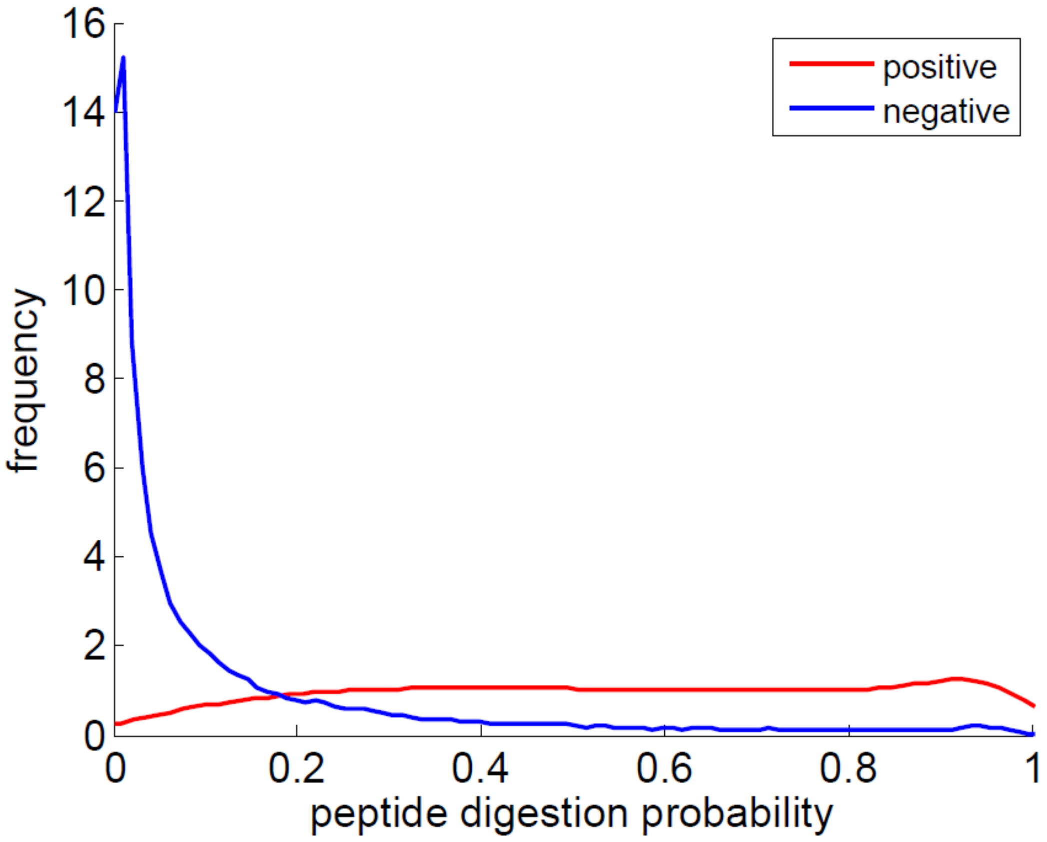


Figure S1. The distribution of the feature “peptide digestion probability” in the training set of the peptide detectability prediction model. The red and blue lines represent the distribution of the feature “peptide digestion probability” of the positive and the negative peptides, respectively.

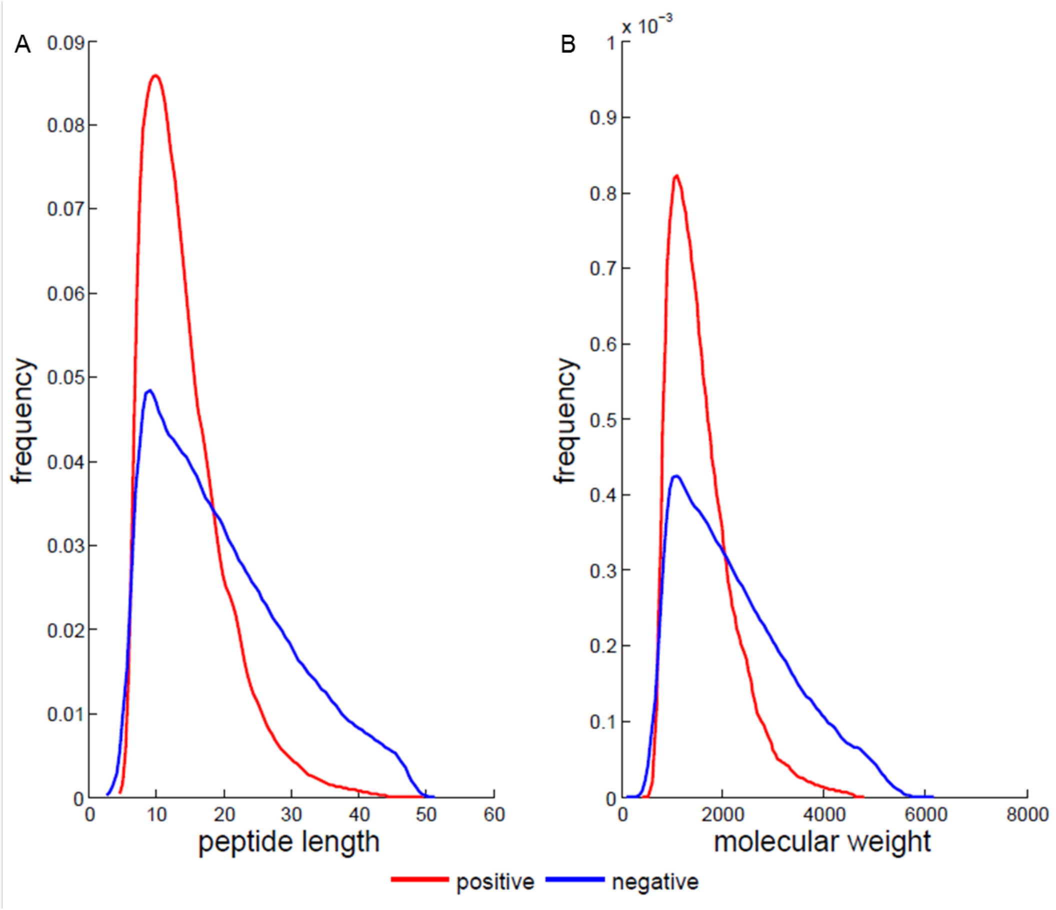


Figure S2. Comparison of the distributions of feature ‘peptide length’ (A) and ‘molecular weight’ (B) on the training set. The red line represents the distribution on the positive peptides and the blue line represents the distribution on the negative peptides.

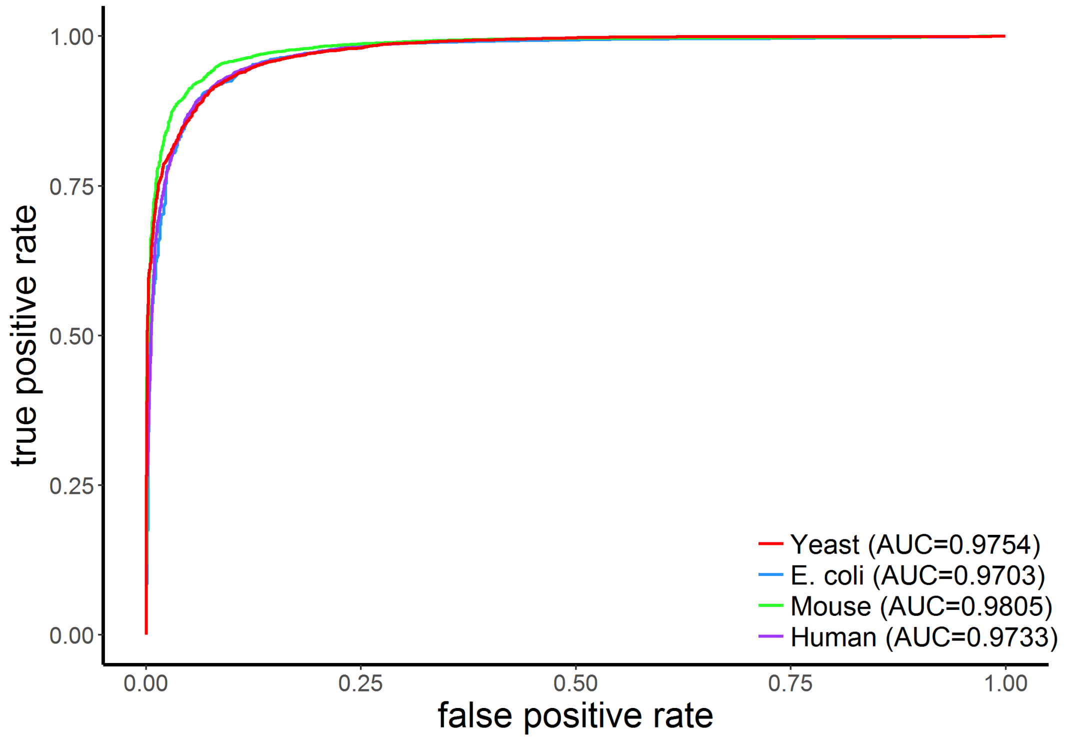


Figure S3. ROC curves demonstrating the performance of cleavage probability model applied to four public datasets. In the yeast dataset, 10-fold cross-validation was performed. In other three datasets, the cleavage probability model was trained on the yeast dataset.

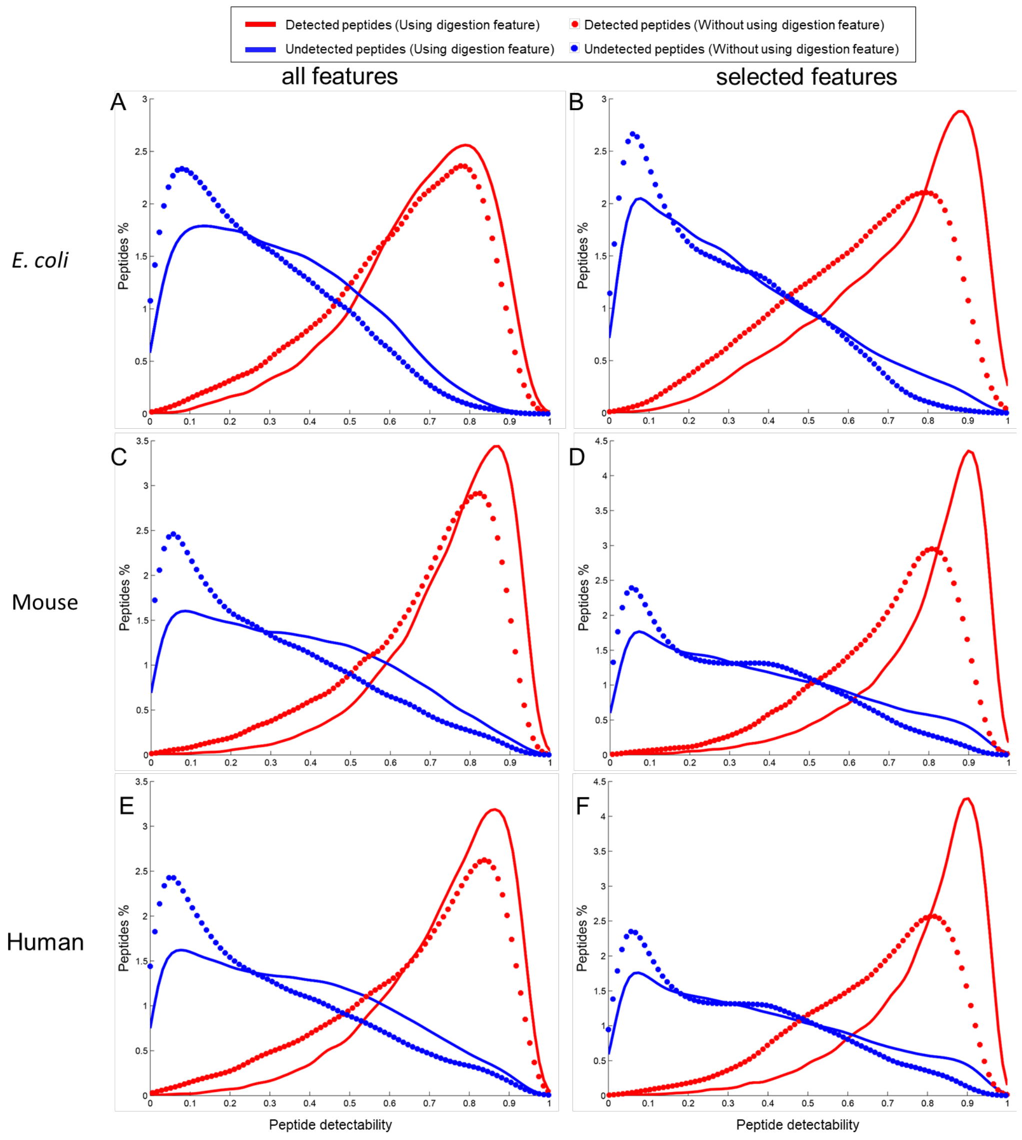


Figure S4 Distributions of predicted peptide detectability for the detected (red) and undetected (blue) peptides of the identified proteins in two cases: using the digestion feature (peptide digestion probability) and without using this feature. (A, C, E) Analyses based on all features for the *E. coli*, mouse and human datasets, respectively. (B, D, F) Analyses based on the selected features for the *E. coli*, mouse and human datasets, respectively.

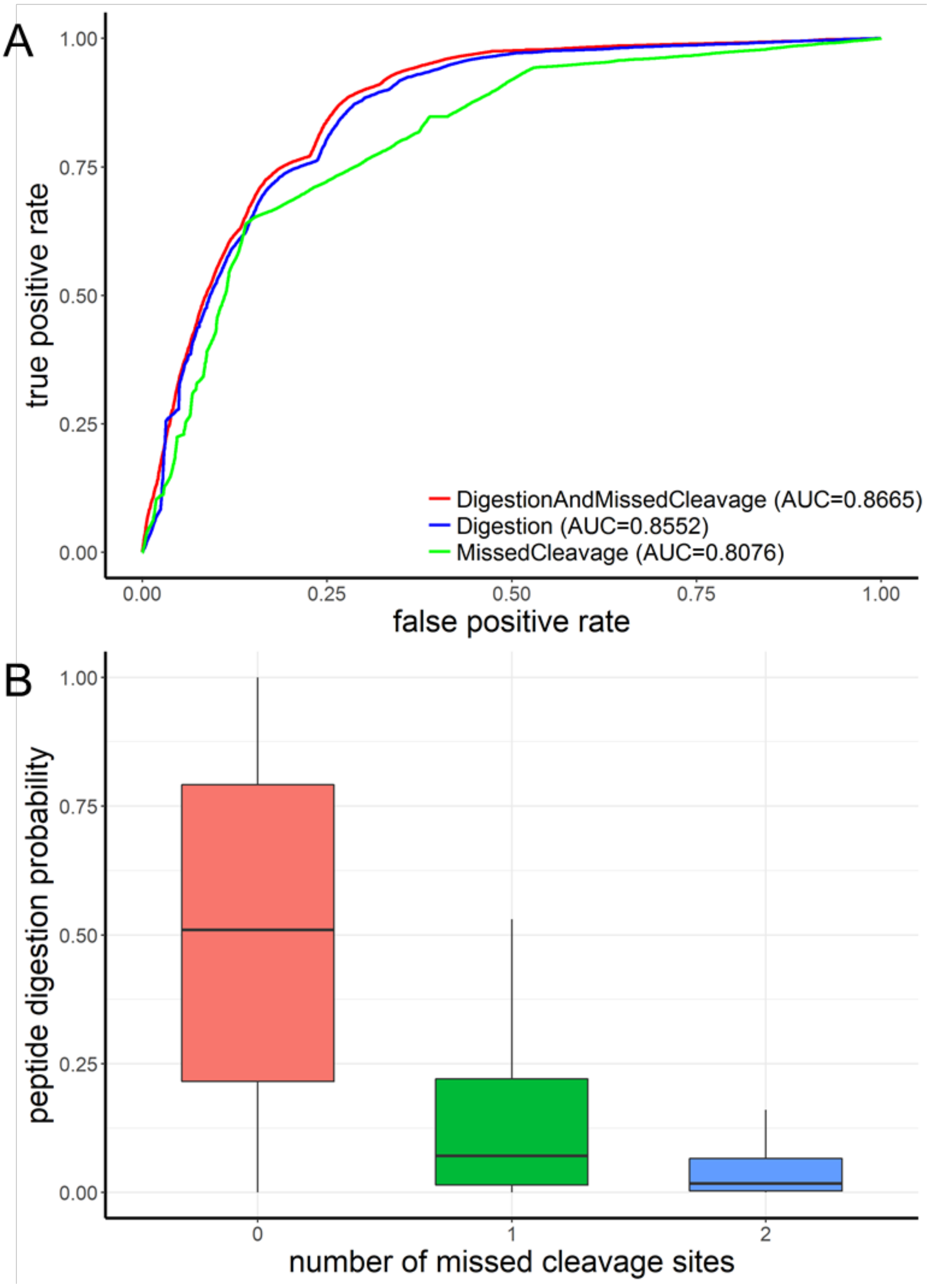


Figure S5. Comparative analysis of peptide digestion probability and the number of missed cleavage sites. (A) ROC curves showing the comparison result of peptide digestion probability and the number of missed cleavage sites. The green, blue and red lines represent the 10-fold cross-validation ROC curves of the models using the number of missed cleavage sites alone, the peptide digestion probability alone and both features, respectively. (B) Boxplots of the predicted digestion probabilities for the peptides with different numbers of missed cleavage sites.

#### Supplement Tables

Table S1. The 588 features used to characterize the peptides in AP3

Table S2. The lists of selected features in three test datasets by mRMR

Table S3. Validation result on MRM assay dataset

(See excel files)
